# Effect of cholesterol on permeability of carbon dioxide across lipid membranes

**DOI:** 10.1101/2020.11.16.384958

**Authors:** M.C. Blosser, J. So, M.S. Madani, N. Malmstadt

## Abstract

Determining the permeability of lipid membranes to gases is important for understanding the biological mechanisms of gas transport. Experiments on model membranes have been used to determine the permeability of lipid bilayers in the absence of proteins. Previous measurements have used a number of different methods and obtained widely varying results. We have developed a microfluidic based microscopy assay that measures the rate of CO_2_ permeation in Giant Unilamellar Vesicles (GUVs), and we report permeability data for the POPC-cholesterol system. We find that cholesterol has a strong effect on permeability; bilayers containing high levels of cholesterol are an order of magnitude less permeable than bilayers without cholesterol, 9.9 ± 1.0 x 10^−4^ cm/s vs. 9.6 ± 1.4 x 10^−3^ cm/s.

**Statement of Significance:** Diffusion of dissolved gasses such as carbon dioxide through cell membranes is an important step in physiological processes. Key to understanding the behavior in cells is the measurement of gas diffusion through model lipid membranes, which isolates the effect of the lipids from other membrane components and allows for control of the composition. Previous measurements have yielded different results for the magnitude of gas transport, and have disagreed on the amount that cholesterol affects transport. The present study presents new data on gas transport across lipid mixtures containing cholesterol, and develops a microfluidic assay for gas transport that will enable further work.

## Introduction

Diffusion of dissolved gasses such as carbon dioxide through cell membranes is an important step in physiological processes, especially for the ability of erythrocytes to clear carbon dioxide. Several proteins, such as aquaporins, the RhAG complex, and AmtB, have been proposed to be gas transporters (1–4). There is currently a debate about the extent to which carbon dioxide transport is dominated by protein transporters or by passive transport through the lipid bilayer itself (5–7). This debate hinges on disagreements about the quantitative measurement of membrane permeability to carbon dioxide. This is compounded by disagreements about the correct theoretical framework for predicting gas transport (8).

Previous measurements of gas transport across lipid membranes have resulted in a broad range of disagreeing values. Recent measurements range from > 3.2 cm/s (9) to ∼0.001 cm/s (10), with additional reports of 0.35 cm/s (11) and .0019 cm/s (12). Itel et al. measured values of .0035 cm/s to >0.16 cm/s for varying amounts of cholesterol (13). Often cited sources for this divergence of measurements are the presence of unstirred layers (USLs) (14), as well as the deadtime involved in mixing solutions.

Traditionally, the permeability of carbon dioxide through lipid membranes has been described with a solubility diffusion model (15, 16). In this model, the membrane acts as a homogeneous hydrophobic slab. The permeability is determined by the partitioning of the solute to the organic phase, the thickness of the slab, and the diffusion coefficient of the species in the membrane environment. This is summarized with

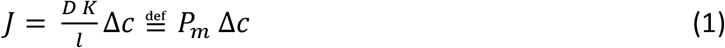

where is *J* the flux of CO_2_ across the membrane, *D* is the diffusion coefficient in the membrane environment, *K* membrane-aqueous partition coefficient of CO_2_, *l* is the membrane thickness, Δ*c* is the difference in CO_2_ concentration across the membrane, and the permeability, *P*_m_, is defined as the ratio of *J* to Δ*c*. This model is often simplified even further by assuming that differences in membrane thickness and diffusion coefficients are small, and so the permeability of a molecule is determined by its partition coefficient between the organic and aqueous phases. This model predicts a permeability on the order of 10 cm/s (9).

In this work, we measure the diffusion of carbon dioxide across membranes composed of POPC and varying amounts of cholesterol. We use an assay combining microfluidics and fluorescence microscopy similar to that in Cama et al. (17), schematically represented in Fig. 1. We use a microfluidic device to rapidly and controllably combine a CO_2_-rich buffer with GUVs. The GUVs contain the pH-sensitive fluorophore HPTS, which decreases in fluorescence as CO_2_ permeates the membrane and acidifies the lumen. This scheme provides for high-throughput measurement while maintaining single-vesicle information. GUVs are an attractive model system for permeability measurements because they are defect and solvent free, and have a large size. Because the characteristic time scale is proportional to the surface to volume ratio, permeation across a 10 µm GUV will take a hundred times longer to equilibrate than the same permeation across a 100 nm large unilamellar vesicle, making fast kinetics significantly easier to measure.

**Figure 1.**
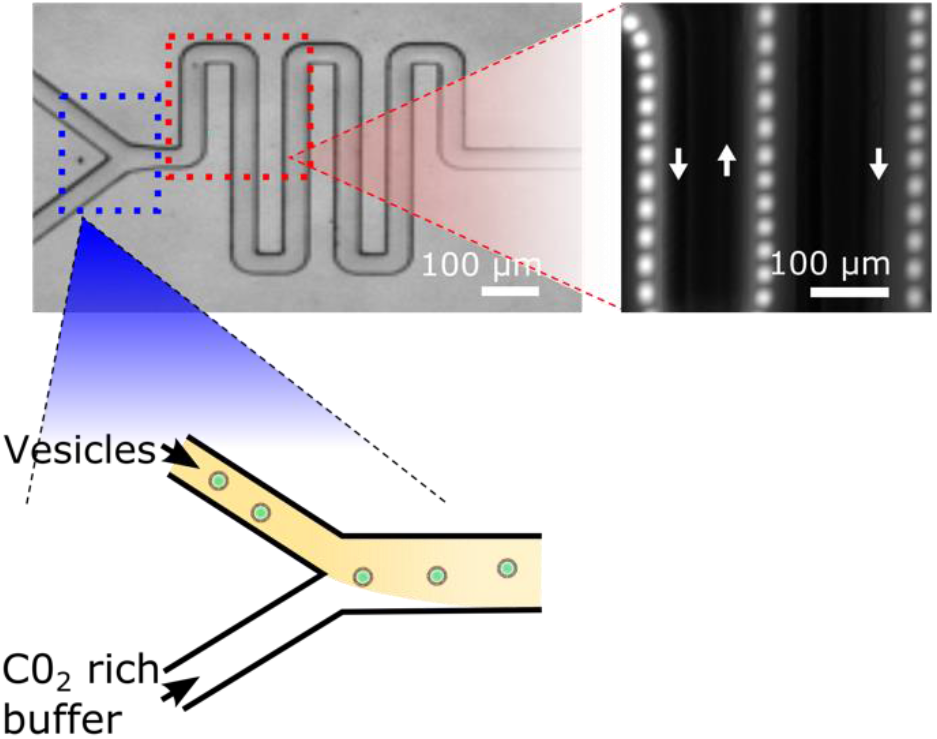
Schematic representation of the experiment. Buffers enriched in CO_2_ and buffer containing vesicles encapsulating pH-sensitive fluorophore mix in a Y-channel. As the solutions mix, CO_2_ diffuses across the lipid membrane, lowering the pH and the fluorescence level. This is imaged as the vesicle traverses the serpentine channel (bright-field micrograph), allowing the same vesicle to be imaged over a longer period of time without immobilizing it. The fluorescence micrograph shows a pseudo-strobe time lapse formed by superimposing micrographs of the same vesicle taken 20 ms apart.

## Materials and Methods

1-palmitoyl-2-oleoyl-glycero-3-phosphocholine (POPC) and 1,2-dipalmitoyl-sn-glycero-3-phosphoethanolamine-N-(lissamine rhodamine B sulfonyl) (rhodamine-DPPE) were obtained from Avanti Polar Lipids (Alabaster, AL). Cholesterol was obtained from Sigma (St. Louis, MO). All lipids were used without further purification and were stored in chloroform at −20°C until use. Pure, 18 mΩ water was obtained from a Milli-Q (MilliporeSigma, Burlington, VT). All other chemicals were obtained from VWR (Radnor, PA).

### Buffers

Vesicles were formed in a buffer consisting of 106 mM NaCl, 2 mM KCl, 1 mM HEPES, 200 mM sucrose, 0.1 mg/mL (∼3.3 µM) carbonic anhydrase, and 0.1 mg/mL (187 µM) of the pH sensitive fluorophore HPTS. The vesicle dilution buffer was composed of 102 mM NaCl, 2 mM KCl, 5 mM HEPES and 200 mM glucose. The CO_2_-rich EQ buffer consisted of 82.8 mM NaCl, 2 mM KCl, 5 mM HEPES, 200 mM glucose, and 33 mM NaHCO_3_. The pH was adjusted to 7.5 before adding the NaHCO_3_. Shortly before an experiment, CO_2_ was bubbled through the solution until the pH reached 7.5. All buffers were checked in an osmometer (Gonotec Osmomat 3000, Berlin, Germany) to ensure they were osmotically balanced.

### Producing vesicles

We produced giant unilamellar vesicles (GUVs) using the method of gel-assisted swelling as described previously (18). Briefly, glass slides were cleaned by sonicating for 30 min in 5 M NaOH, then 30 minutes in methanol, and then rinsed thoroughly. They were plasma treated for 3 minutes (Harrick Plasma, Ithaca, NY) immediately before being coated with agarose. 25 µL of 2% w/v molten ultra-low gelling temperature agarose was pipetted onto one slide, then another slide was placed on top, and the two slides were slid past each other. The slides were allowed to gel at room temperature overnight. 15 µL of a lipid solution at 2 mg/mL in chloroform was added to a slide, and immediately dried under a stream of nitrogen to remove the chloroform and create an even film of lipids. Lipid solutions always included 0.2 mol% rhodamine-DPPE for imaging the membranes. These films were put under 400 µL of vesicle formation buffer and allowed to rehydrate for 1 hour. Prior to harvesting, the vesicles were gently rinsed ten times by slowly removing 200 µL of buffer and replacing it with 200 µL of dilution buffer. During this process the majority of GUVs remained adhered to agarose hydrogel. The solution was then agitated by gently pipetting parallel to the surface and harvested. A sample from each vesicle batch was imaged in the rhodamine channel to ensure vesicles had formed correctly. For permeability experiments, the vesicle solution was diluted in the dilution buffer ∼10:1 to ensure that images contained multiple vesicles only rarely. Vesicles were always formed the same day as experiments.

### Microfluidics

We produced PDMS (polydimethylsiloxane) microfluidic channels with rectangular cross sections. The channels were produced by replica molding from a mask consisting of a Y junction followed by a serpentine channel (Fig. 1). A glass coverslip (VWR) that was cleaned as above was plasma treated together with the PDMS channel for 30 s, after which they were immediately bonded to one another. Before use, the channel was incubated with 1 mg/mL bovine serum albumin for 30 min. to passivate surfaces in order to minimize GUV rupture. The vesicle solution and CO_2_-enriched solutions were loaded into 500 µL Hamilton gas-tight pipettes (Reno, NV) connected to 0.02 inch ID Tygon tubing (Saint Gobain, France). Flow rate was controlled by a Fusion 200 syringe pump (Chemyx, Stafford, TX) at a rate of 0.4 µL/min.

### Imaging

Flowing vesicles were imaged in epifluorescence as in (18). Briefly, we used an Axio Observer Z1 miscroscope with an EC Plan-Neofluar 40× objective and a Colibri 2 LED Illumination System with a 120 V LED (Zeiss, Germany). Images were captured using a Hamamatsu CMOS camera (Hamamatsu, Japan). Exposure time was 10 ms, with a rolling shutter. Images were binned 4×4 to increase signal and reduce storage requirements for long acquisitions with a sparse population of vesicles.

Quantitative fluorescence levels were extracted using a custom python script. Images were first corrected for dark noise and flat field. Frames potentially containing vesicles were identified by subtracting the median projection of the entire stack, and then comparing the residual maximum intensity to a manually set threshold. Frames identified in this way were subjected to a more refined background subtraction, subtracting the median of 5 frames before and after the frame of interest. The resulting image was subjected to an Otsu filter to identify vesicles. Because of significant motional blurring, vesicle size was determined by from the dimension perpendicular to the direction of motion. The maximum fluorescence was determined from the median value of the brightest 10 pixels. We chose this to reduce error introduced by inaccuracies in identifying vesicle edges while avoiding large fluctuations in single pixels. Events were then manually chosen by selecting vesicles which appeared round (as opposed to aggregates) and for which data was successfully recorded for all three passes across the field of view.

## Results and Analysis

Membrane permeability to CO2 was measured for POPC membranes with a range of cholesterol concentrations from 0-60 mol%. In all cases, the fluorescence decay was well described by an exponential,

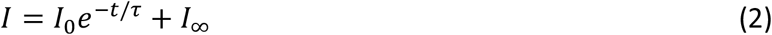

as seen in Fig. 2. This agrees with expectations that the change in fluorescence is dominated by a single process.

**Figure 2.**
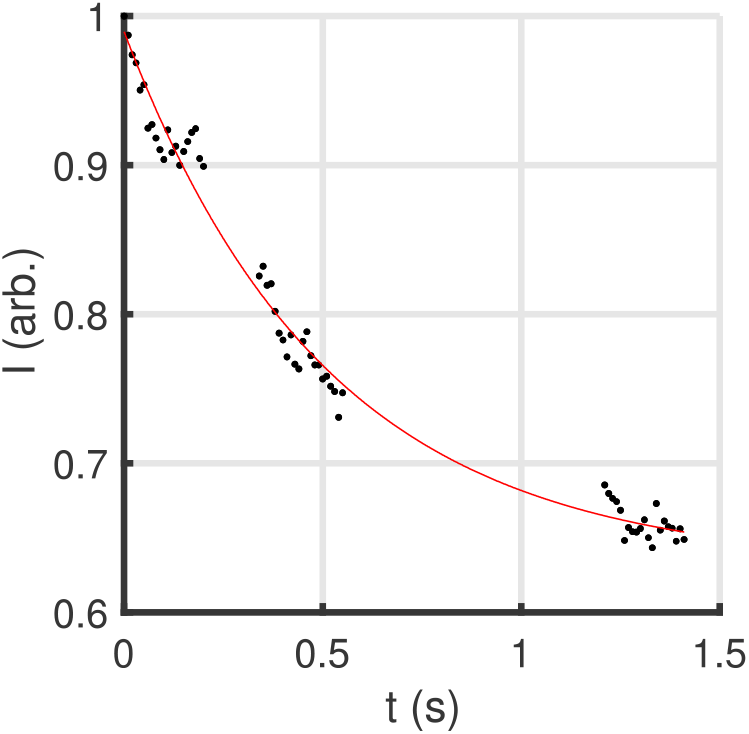
Exemplary data from a vesicle with composition 40-60 POPC-chol. The fluorescence has been normalized to the first point. Data points are derived from each frame of the acquisition. The breaks in the data arise from the vesicle leaving the field of view as it traverses the channel. The red line is a fit to Eq. 2, with a 1/τ = 1.58 s^-1^ and *I*_∞_ = 0.63.

A naïve model of transport of species with permeability *P*_m_ obtained by integrating equation 1 is:

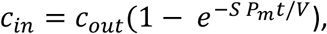

where *c*_*in/out*_ is the concentration inside/outside the GUV, *S* is the surface area, and *V* is the volume. Since a substantial fraction of the CO_2_ will be hydrated to form carbonate, the concentration inside will grow more slowly than equation 2 predicts, so that

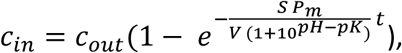

where *pK* is the pKa of CO_2_. However, we do not directly measure the internal concentration of CO_2_; the measured signal is proportional to the concentration of deprotonated HPTS. We also cannot assume that the pH of the system will be constant. To take into account both the finite buffer capacity of the system, and the finite response time of the other components of the system, we numerically solved a system of coupled rate equations describing protonation/deprotonation of each of the components, as in (19). The rate constants of each component were obtained from literature values. The details of these equations are in the SI. To enable fitting a large number of curves, each with a different surface to volume ratio, we solved the system of equations for a variety of permeabilities to produce simulated data, assuming that the measured fluorescence is proportional to the concentration of unprotonated HPTS. From this simulated data with specified permeability we extracted the effective time constant 1/τ and final intensity, *I*_∞_ by fitting to equation 2. The numerical relationship (Fig. S2) was then used to interpolate permeabilities from a measured 1/τ. Using this relationship, the 1/τ of each vesicle was separately translated to a value of *P*_m_. The error presented is the standard error of the mean arising from the variance in the data for a given set of conditions. To account for the finite mixing time, an offset of 0.06 s was introduced to the modeled data based on observations of the time vesicles required to travel from the Y-junction to the serpentine.

We measured the permeability of a range of cholesterol concentrations. The results are shown in Fig. 3. We find an order of magnitude decrease in *P*_m_ between cholesterol-free membranes and membranes with 60 mol% cholesterol, which is near the cholesterol solubility limit of 66 mol% (20). This dependence on cholesterol gives further confirmation that the measured permeability is a measurement of properties of the lipid bilayers themselves. Specifically, it implies that neither USLs nor slow mixing of buffers are dominating the barrier to diffusion.

**Figure 3.**
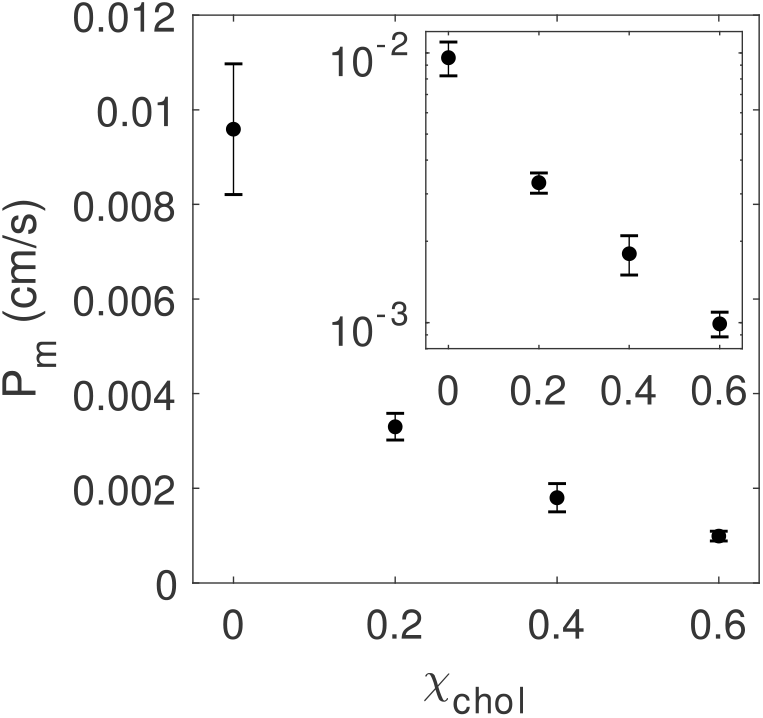
Permeability of CO_2_ as a function of cholesterol mol fraction χ_chol_. Error bars represent the standard error of the mean. N = 22, 20, 29 and 21 from low to high cholesterol respectively. The inset shows the same data on a linear-logarithmic scale to better represent the relative uncertainties.

To ensure that the conversion of CO2 to carbonic acid was not the rate-limiting step, we varied the amount of carbonic anhydrase (Fig. S3). We found that increasing the concentration by a factor of 1.5 had no significant effect on the measured permeability. This is consistent with 0.1 mg/ml being saturating for our conditions. We also found that completely removing the carbonic anhydrase results in kinetics too slow for our experimental setup to measure. This confirms that conversion of CO_2_ to carbonic acid is a necessary step in the measurement of CO_2_ permeability, consistent with the chemical pathway in our model.

To validate that our results were not an artifact of finite buffer capacity, we varied both the external concentration of CO_2_ and the concentration of the buffer HEPES encapsulated in the vesicles. The results are summarized Fig. S4 and Fig. S5. Decreasing the external concentration of CO_2_ results in no significant change in the time constant of the fluorescent decay 1/τ, but a significant decrease in the size of the fluorescence drop, (*I*_0_ − *I*_∞_)/*I*_0_. Similarly, increasing the buffer capacity did not significantly change 1/τ, but decreased the size of the fluorescence drop. In both cases, the implication is confirmation that the change in fluorescence is due to a CO_2_-induced pH change, and that the buffer dynamics of the system are not dominating the reaction kinetics.

## Conclusion

In this paper, we developed a microfluidic based fluorescence assay to measure permeabilities of CO_2_ across lipid membranes and used this method to determine the effect of cholesterol concentration on CO_2_ permeability. This technique relies on transient gradients, rather than steady state measurements that are more sensitive to the details of unstirred layers. The use of micron-scale vesicles allows the characteristic equilibration time to be resolved using conventional fluorescence microscopy techniques. When combined with techniques for reconstituting proteins into GUVs, this will allow the *in vitro* measurement of permeability through membrane channels such as aquaporins.

The substantial effect of cholesterol seen in this work is consistent with membranes being a barrier to diffusion of gasses not well modeled by a single hydrophobic layer, similar to arguments made in recent simulation work (8). The effect of cholesterol agrees in direction, but not magnitude, with previous work by Itel et al. (13). Simulation work by Hub et al. also reported a strong effect of cholesterol on CO_2_ permeability (21). Interestingly, they saw only small changes at low levels of cholesterol, increasing as the cholesterol content increases. In contrast, we measure a large change in permeability between 0 and 20 mol% cholesterol, with the absolute change in permeability decreasing with increasing cholesterol content (the work by Itel et al. was not sensitive at low concentrations of cholesterol).

Cholesterol is known to increase the thickness of and decrease lateral diffusion in lipid bilayers. However, neither of these effects explains our results in the solubility diffusion framework. Hung et al. measured the thickness of POPC bilayers changing only for 5.3 nm at 0 mol% chol to 5.8 nm at 38 mol% chol (22). This would predict a 10% change in the permeability over this range, while we observe close to an order of magnitude difference. Similarly, the lateral diffusion of lipids decreases with increasing cholesterol, suggesting an increase in viscosity. Filipov et al. measured a ∼40% decrease in diffusion coefficient in 50 mol% POPC bilayers compared with pure POPC (23). In both cases, the observed changes are approximately linear, unlike the behavior observed in this work.

## Author Contributions

MCB, JS, and NM designed experiments; MCB, JS, and MSM performed experiments; MCB and MSM analyzed data; and MCB and NM wrote the manuscript.

## Acknowledgements

This work was supported by Office of Naval Research Subcontract FP00209340. The authors thank Sepehr Maktabi for fabricating the channel mold, and Walter Boron for helpful conversations.

